# Label-free toehold mediated strand displacement on 3D printed hybrid paper-polymer platform for protein sensing

**DOI:** 10.64898/2026.03.27.714923

**Authors:** P. Ngaju, D. Kakadiya, Sorosh Abdollahi, K. Kim, R. Pandey

## Abstract

A programmable 4-input cascade DNA logic gate utilizing toehold mediated strand displacement (TMSD) was implemented on a 3D printed hybrid paper-polymer vertical flow device (3D HPVF) for on/off sensitive and specific fluorescence detection of platelet derived growth factor BB (PDGF BB). Polypropylene was 3D printed directly on paper and thermally cured to create micro paper analytical devices (µPADs). The 3D HPVF comprised of three layers of µPADs enclosed in a casing that clamped each µPAD securely to ensure seamless and efficient wicking between layers. In the presence of PDGF BB, a partially complementary strand to a PDGF B aptamer (PDGF B Apt), cApt, was liberated from a PDGF B Apt/cApt duplex in solution. The solution was then deposited on the 3D HPVF with a dimeric g-quadruplex hairpin. The 4-nucleotide toehold region on the cApt started the hybridization reaction with the dimeric g-quadruplex hairpin (dGH) opening it up allowing formation of a dimeric g-quadruplex structure that binds with thioflavin T (ThT) with enhanced fluorescence intensity at room temperature. The 3D HPVF exhibits a pico molar range of detection from 10pM to 100pM with a 10pM limit of detection (LOD) for PDGF BB concentrations relevant for pregnant women predisposed to early-onset preeclampsia with clear differentiation when compared to similarly competing analytes PDGF AA and AB.

## Introduction

PDGFs belong to a family of dimeric glycoprotein growth factors composed of A, B, C and D chains forming five isoforms (PDGF AA, AB, BB, CC, DD) linked by disulfide bonds, these ligands bind to two structurally related tyrosine kinase receptors PDGF Rα and PDGF Rβ, with PDGF BB as the most bioactive isoform^1,2^. In early-onset preeclampsia (less than 34 weeks gestation), placental insufficiency as a result of shallow trophoblast invasion and defective spiral artery remodelling, PDGF BB plays a pathological role in vascular maladaptation^3^. PDGF BB contributes to decidual blood vessel pathology, driving excessive pericyte/smooth muscle proliferation, impairing spiral artery dilation needed for uteroplacental perfusion^4^. Based on the literature, fluorescence paper-based assays are limited. Most PDGF BB fluorescence strand displacement aptasensors in the literature incorporate fluorophore labelled nucleic acids and are implemented in solution. Liu et al. presented findings on a dynamic DNA assembly mediated by binding-induced DNA strand displacement within a homogenous solution assay wherein a PDGF B aptamer was split between two DNA probes that form a complex. PDGF BB binding brings the complexes together, releasing an output strand that initiates a toehold mediated strand displacement that liberates a fluorophore from a fluorophore-quencher duplex, turning fluorescence on^5^. Paper microfluidic fluorescent aptasensors have been implemented for SARS-CoV-2 spike protein, matrix metalloproteinase-2 (MMP-2), carcinoembryonic antigen (CEA) and other protein targets but not for PDGF BB^6,7^. For instance, Basabe-Desmonts et al. developed a paper based microfluidic platform for single step detection of mesenchymal stromal cells secreted vascular endothelial growth factor (VEGF)^8^. This device used a three-part structure switching aptamer on paper with an affinity to VEGF which provided a quantifiable fluorescent signal by displacement of a quencher when VEGF was successfully bound^8^.

To the best of our knowledge, label free strand displacement paper-based biosensors towards protein sensing are yet to be explored. The current gold standard for early stage preeclampsia screening focuses on angiogenic imbalance biomarkers, soluble fms-like tyrosine kinase 1 (sFlt-1)/placental growth factor (PlGF) ratio, and are currently immunoassays run in the laboratory on commercial analysers such as those manufactured by Roche Diagnostics, Siemens and Thermofisher scientific^9–11^. Despite the high performance of commercial immunoassay analysers, they require centralized laboratories, skilled technicians, controlled temperature and regular maintenance rendering them unsuitable for rural and resource constrained settings that require accurate isothermal testing at the point of care. Furthermore the reagent cost per test for the patient including instrument overhead is incredibly high, sometimes costing thousands of dollars^12^. Paper-based diagnostic tests for preeclampsia reported in the literature have detected circulating plasma proteins FKBPL and CD44, and GlyFn implemented on lateral flow tests including colorimetric urine protein aggregation tests such as Congo red dot (CRD) and dipstick urinalysis^13–15^. The FKBPL and CD44, and GlyFn lateral flow tests utilize antibodies as the capture bioreceptor whereas CRD and dipstick tests detect misfolded and damaged proteins present in the urine of preeclamptic patients typically after 20 weeks. PDGF BB offers several advantages over the CRD tests, primarily in specificity, mechanistic insight and stage prediction. CRD detects non-specific protein aggregates (congophilia) which may have overlap with proteinuria (late stage preeclampsia), whereas PDGF BB is linked specifically to vascular remodelling pathology during early onset^16^. Furthermore, nucleic acids such as aptamers offer critical advantages over antibody paper-based tests. Aptamers are known for their chemical stability allowing them to withstand room temperature storage once immobilized on paper, high sensitivity and selectivity to their target analyte, minimal batch to batch variation and considerably lower production costs in comparison to antibodies^17^.

Based on this background, this work proposes PDGF B Apt/PDGF BB binding initiated TMSD, an isothermal programmable DNA hybridization reaction wherein a short single-stranded ‘toehold’ region on an initiator strand provides an initial binding site to open dGH, providing low background, tuneable kinetics and high specificity ideal for biosensing. TMSD is implemented in solution for verification and finally on a 3D HPVF for sensitive and selective detection of PDGF BB using a dimeric g-quadruplex with ThT as a fluorescent probe. Dimeric g-quadruplexes dramatically amplify ThT by more than 100-fold when compared to their monomeric counterparts, enabling detection in the nano molar to picomolar range due to multiple ThT binding sites on the dimeric interface, exhibition of stable fluorescence, coupled with compatibility to paper^18,19^. The 3D HPVF utilizes a low sample volume of 20 µl (10 µl each for the test and control) ideal for typically low abundance biomarkers such as PDGF BB and exhibits repeatable sensitivity between 10pM to 100pM with selectivity against PDGF AA and PDGF AB. The proposed 3D HPVF for PDGF BB is mechanistic and angiogenic, hence appropriate for use as an early onset screening tool alongside existing clinical care guidelines for preeclampsia.

## Materials and Methods

### Reagents and Materials

Ethylenediaminetetraacetic acid ((HO_2_CCH_2_)_2_NCH_2_CH_2_N(CH_2_CO_2_H)_2_) (EDTA), IDTE (1xTE) buffer (10mM Tris, 0.1mM EDTA) at pH 7.5, 1xTBE buffer (50mM Tris-H_3_BO_3_, 2mM EDTA.2Na, pH 8.5). Tris hydrochloride(tris-HCl), Thioflavin T (ThT), ammonium persulfate (APS), ethidium bromide, N,N,N’,N’-Tetramethylethylenediamine (TEMED), 30% Acrylamide/Bis-acrylamide, sodium chloride (NaCl), ultrapure DNase/RNase-free distilled (MilliQ) water were all purchased from Millipore Sigma (Vermont, US). Gel red nucleic acid stain, recombinant human platelet derived growth factor (PDGF) BB, PDGF AB, PDGF AA, proteins were obtained from ThermoFisher Scientific (Massachusetts, US). Deionized water (DI) used in all experiments had a resistivity of 18.2 MΩ . cm at 25 °C. All DNA sequences used are unique to this study with the exception of the PDGF B aptamer which was obtained from the literature^20^. All DNA sequences were purchased from Integrated DNA Technologies (Iowa, US) and are listed in ***Table S1***. Polypropylene (PP) and polylactic acid (PLA+) filaments were obtained from Shop3D (Mississauga, ON). Whatman Grade 1 cellulose chromatography paper with a thickness of 0.18mm and a linear flow rate of 130mm/30min was purchased from Cytiva Life Sciences (Massachusetts, US).

### Fabrication of 3D hybrid paper-polymer micro analytical platform

A vertical flow device with three layers and two reaction pads with a diameter of 2mm was designed using SolidWorks Academic Version (Autodesk). An FDM 3D printer (Qidi Tech X-Plus II) (China) was used to 3D print hydrophobic barriers using polypropylene directly on chromatography paper. Thereafter the paper device was thermally cured in a convection oven for 27 minutes and allowed to cool for 5 minutes before using in experiments. A casing was designed to house the 3D hybrid paper-polymer µPAD device (3DH-µPAD)using SolidWorks Academic Version (Autodesk) and 3D printed using PLA.

### Affinity of PDGF B Aptamer

12% native polyacrylamide gel electrophoresis (PAGE) was used to demonstrate the affinity of the PDGF B aptamer to PDGF BB. 10 µM of aptamer was diluted in 1X annealing buffer from a 100 µM stock solution in TE buffer, heated at 90-95°C for 5 minutes and cooled for 1 hour at room temperature. The 1X annealing buffer was 10mM NaCl, 50mM Tris-HCl and 1mM EDTA prepared from a 10X stock solution of 100mM NaCl, 500mM Tris-HCl and 10mM EDTA. PDGF BB was reconstituted with sterile water as per the manufacturer’s specifications 5µl of the aptamer was incubated with 5µl of PDGF BB at four different concentrations at 37°C in the dark, specifically 10µM, 5µM, 3µM and 1µM for 30 minutes. The gel was run in 1xTBE buffer for 60 minutes at a voltage of 100 V. After electrophoresis, the gel was stained in ethidium bromide for 30 minutes and then imaged using a BioRad ultraviolet (UV) imaging system. The free PDGF B aptamer was compared to the aptamer/protein complexes to determine aptamer-protein binding.

### Gel electrophoresis experiments to validate TMSD assay

Oligonucleotides were re-constituted in TE buffer to make 100µM stock solutions and then subsequently diluted using a 1X annealing buffer. The PDGF B Apt, initiator strand complementary to the PDGF B Apt (cApt), dGH sequences used within this study were diluted to 2µM solutions using the 1X annealing buffer. To hybridize the PDGF B Apt to the cApt forming a duplex, oligonucleotides were heated up to 95°C for 5 minutes on a heating block and thereafter slowly cooled to 22 °C for 1 hour. The dGH was formed by heating up to 95°C for 5 minutes on a heating block and thereafter slowly cooled to 22 °C for 1 hour. After hybridization the PDGF B Apt/cApt complex was incubated with PDGF BB protein for 30 minutes at 22°C and thereafter the dGH for 1 hour at 22°C. Controls and interactions were verified using 12% PAGE. Native polyacrylamide gel in 1xTBE buffer was run for 60 minutes at a voltage of 100 V. After electrophoresis, the gel was stained in ethidium bromide for 30 minutes and then imaged using a BioRad UV imaging system.

### Study on the effect of concentration on fluorescence intensity

A solution assay was performed to study the effect of DNA concentration on overall fluorescence intensity. Samples were prepared in triplicate and the concentrations used were 75, 100, 125, 150 and 175nM. Fluorescence intensity of the controls PDGF B Apt, PDGF B Apt/cApt, PDGF B Apt/cApt/PDGF BB were compared with the full assay PDGF B Apt/cApt/PDGF BB/dGH. The concentration of PDGF BB used for the experiment was 1nM. Fluorescence intensity was read using a BMG Clariostar microplate reader, with an excitation wavelength of 425nm, an emission wavelength of 482nm and enhanced dynamic range.

### Study on the effect of incubation time and temperature on fluorescence intensity

A 3 x 5 factorial design experiment was conducted to determine the effect of temperature at three levels (4 °C, 22 °C and 37°C) and incubation time at five levels (30min, 1 hr, 2hrs, 3 hrs and 4hrs) on the fluorescence intensity of the full assay PDGF B Apt/cApt/PDGF BB/dGH. The concentration of PDGF BB used for this experiment was 1nM. PDGF B Apt was hybridized with cApt, incubated with PDGF BB for 30 minutes and then with dGH for 1 hour as detailed in the protocol described in 4.2.4. Fluorescence intensity was read using a BMG Clariostar microplate reader, with an excitation wavelength of 425nm, an emission wavelength of 482nm and enhanced dynamic range.

### Assay validation on 3D printed hybrid paper-polymer vertical flow device

10 µl of 5 µM of Thioflavin T was deposited on the sensing regions of the 3DH-µPAD and allowed to dry for 24 hours. The dGH was deposited on the test region and the cApt on the control region. The sample protein was incubated with PDGF B Apt/cDNA in solution and thereafter 10 µl deposited on the test region and another 10 µl deposited on the control reaction pad. The assay was run for 30 minutes and thereafter allowed to completely dry before capturing images of the test and control regions using a Nikon Eclipse Ti-2 fluorescence microscope with the fluorescence filter set to ‘GFP’ with an excitation wavelength of 446-486nm and emission wavelength range of 500-550nm. Image J Fiji licensed under the GNU public licence version 2 was used analyze the intensity of the fluorescence signal to quantify the degree of fluorescence in each image. The experiment was carried out in triplicate (n=3 discrete devices for each concentration).

### Statistical Analysis

All measurements obtained from the experiments were plotted to show mean ± standard deviation. Statistical analysis of the data was performed in GraphPad Prism software version 10.5.0. One-way and two-way ANOVA were used as the statistical tests considering a 95% confidence interval with Tukey’s and Sidak’s multiple comparisons tests. Statistical comparisons between groups are described within the figure captions and each experiment was conducted with a minimum of n=3 samples.

## Results and Discussion

### Toehold mediated strand displacement assay for PDGF BB detection

A TMSD assay was designed for PDGF BB detection wherein the binding event of the PDGF B aptamer (A) and PDGF BB (B) initiated the displacement of an initiator strand (F) with a toehold region that unlocks a dimeric g-quadruplex hairpin (H) (***Figure 1***). The PDGF B aptamer (B) is hybridized partially to an initiator strand (F) such that it has optimal stability, however, not so stable such that PDGF BB binding to A cannot displace the F to initiate the reaction. Base pairing was carefully designed to balance AT and GC base pair content. The ends of the A and B duplex were ‘clamped’ with GC clamps for 3 out of the 4 base pairs at the 5’ and 3’ ends (***Table S1***). A was modified such that its nucleotides were extended at the 5’ end to restrict access to the toehold region (D) to prevent undesired ‘leak’ reactions, ensuring not to alter the core binding region to ensure the affinity of A to B was retained. The toehold region on F is complementary to a toehold region (G) sequestered within a dimeric g-quadruplex hairpin (H). The hybridization of the initiator strand (F) with its complementary region (G) on H is set in motion by the toehold (E) which acts as a kinetic ‘latch’ comprising of a four-nucleotide sequence. This event disrupts the hydrogen bonds between the intramolecular base pairs of the dimeric g-quadruplex hairpin, fully exposing H which subsequently forms a dimeric anti-parallel g-quadruplex structure (I). Dimeric g-quadruplexes have several advantages over their monomeric counterparts when used as fluorescent probes, particularly for dyes such as ThT^21^. G-quadruplexes are known to bind to thioflavin T with enhanced fluorescence intensity in the presence of monovalent cations through end-stacking or groove binding at the terminal G-tetrads with π-π stacking interactions involving exposed guanine bases^22–26^. Dimeric g-quadruplexes provide significantly higher fluorescence and quantum yields due to binding regions between units, allowing the efficient stacking of dyes with improved mobility restriction^25^. I binds with ThT with enhanced fluorescence intensity in the presence of 150mM Na+ as demonstrated in ***Figure 1***.

**Figure 1:**
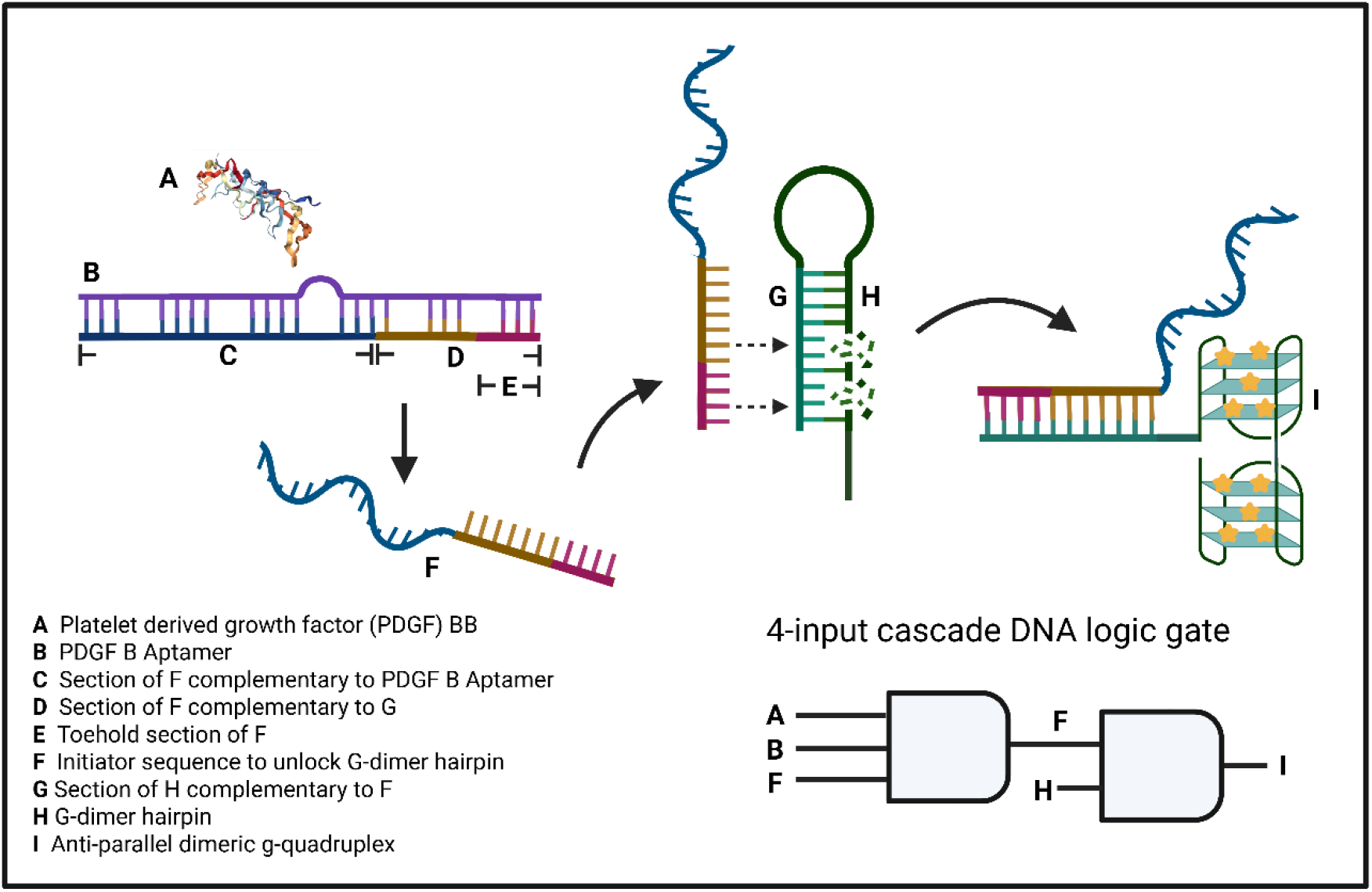
Schematic of the toehold strand displacement assay for detection of PDGF BB. The PDGF B Apt (B) is hybridized to a cApt (F) which is displaced when PDGF BB binds to B. F contains a toehold section that initiates the kinetic reaction of F hybridizing to G which opened the dGH (H) allowing it to form an anti-parallel dimeric g-quadruplex structure (I). This assay is modeled using 4 input cascade DNA logic where a 3-input AND gate receives 3 inputs, A (PDGFBB), B(PDGF B Apt) and F (cApt) with an output that is ‘1’ only when all inputs are 1, this output feeds into the input of a second AND gate whose inputs are F and H, both inputs would have to be ‘1’ in order to unlock the dimeric g-quadruplex sequence to achieve the g-quadruplex structure I.

The entire assay is modelled using a 4-input cascade DNA logic gate (***Figure 1***) that comprises of 3 input and 2 input logic gates as shown in ***Figure 2***. The 3-input logic gate has 3 inputs, PDGF BB protein (A), PDGF B Apt (B), and cApt (F), where B is hybridized with F. The 3-input AND gate can only have a high ‘1’ if all input bits are high ‘1’. The output of the 3 input AND gate is the displaced initiator strand F when A,B and F are all ‘1’ (***Figure 2A***). This output is the one of the inputs for the subsequent 2-input AND gate with the second input as dGH (H). When F hybridized with H, the dimeric g-quadruplex structure is allowed to form, similarly, both F and H must be ‘1’ for I to form (***Figure 2***). To predict the formation of the dimeric g-quadruplex at room temperature (22°C) based on the hybridization event of the initiator sequence to the hairpin, NUPACK software was used as shown in ***Figure 3***. The initiator sequence (cApt) has a Gibbs free energy of -7.96kcal/mol in 150mM Na+ and 2mM Mg at room temperature (***Figure 3A***). When cApt which contains the 4-nucleotide toehold region hybridizes with dGH, there is a significant drop in Gibbs free energy to -32.59kcal/mol, which indicates formation of hydrogen bonds to form a double stranded complex (***Figure 3B***). Hybridization of cApt with its complementary region on dGH exposes the dimeric g-quadruplex nucleotides, allowing the dimeric g-quadruplex structure to form. These results mean that the hybridization event between cApt and dGH is highly probable with a thermodynamically favourable and spontaneous kinetic reaction within the given conditions. This event would allow the exposure of the g-rich sequence responsible for formation of the dimeric g-quadruplex anti-parallel structure which is the sensing probe for the assay.

**Figure 2:**
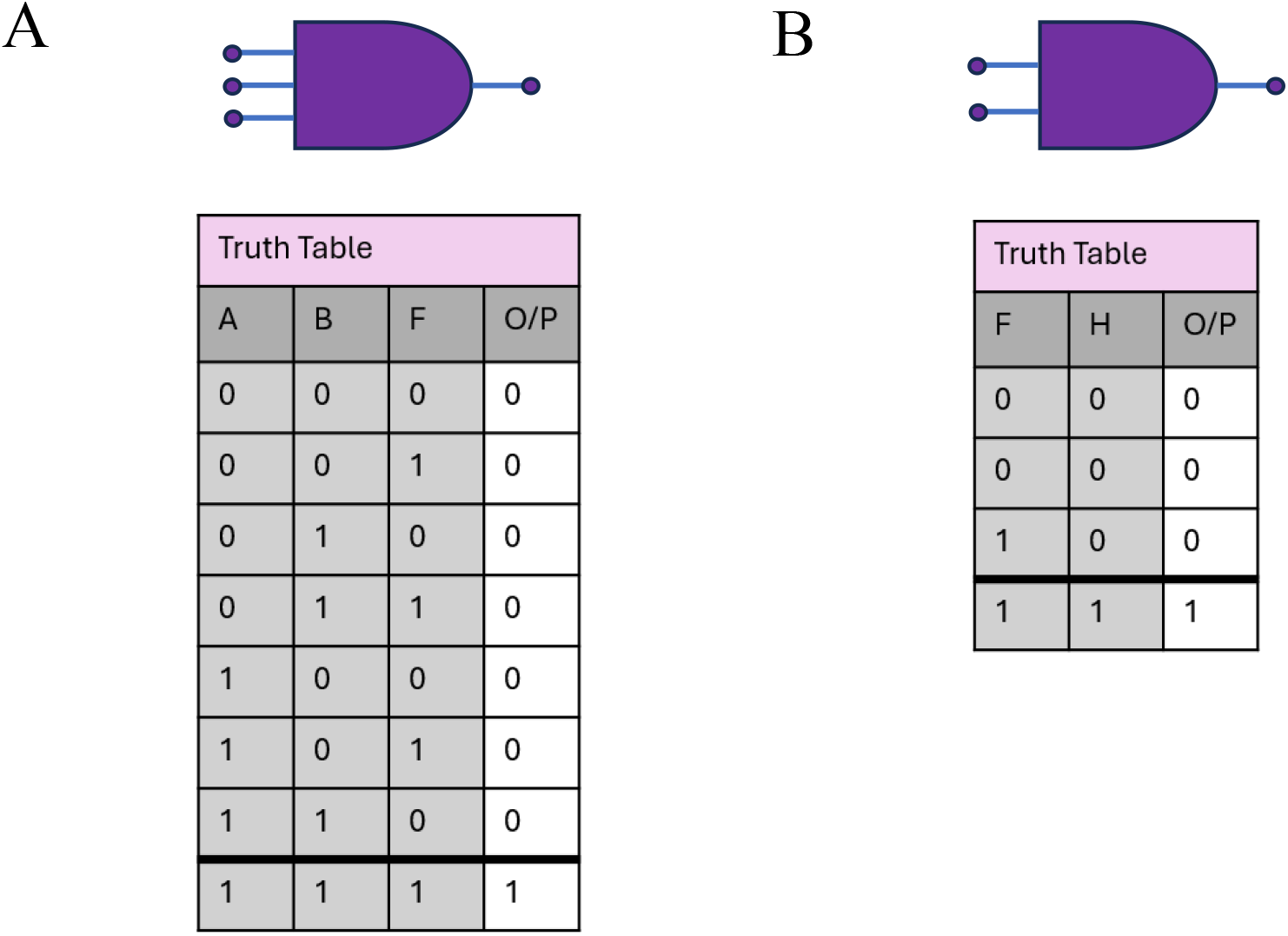
3-input and 2-input AND gates used to model 4-input cascade DNA molecular logic gate for the TMSD assay. **A**. 3-input AND gate with PDGF BB (A) and Apt (B) and cApt (F) as inputs, the output (O/P) bit which is the displaced strand (F) is high’1’ only when all inputs are high meaning A would have to bind to B/F complex to release F. **B**. 2-input AND gate has 2 inputs namely F from the 3-input AND gate and H, F has a toehold at the 3’ end and hybridizes with H at the 5’ end exposing the dimeric g-quadruplex sequence to form the O/P (I) which is high only if F and H are ‘1’.

**Figure 3:**
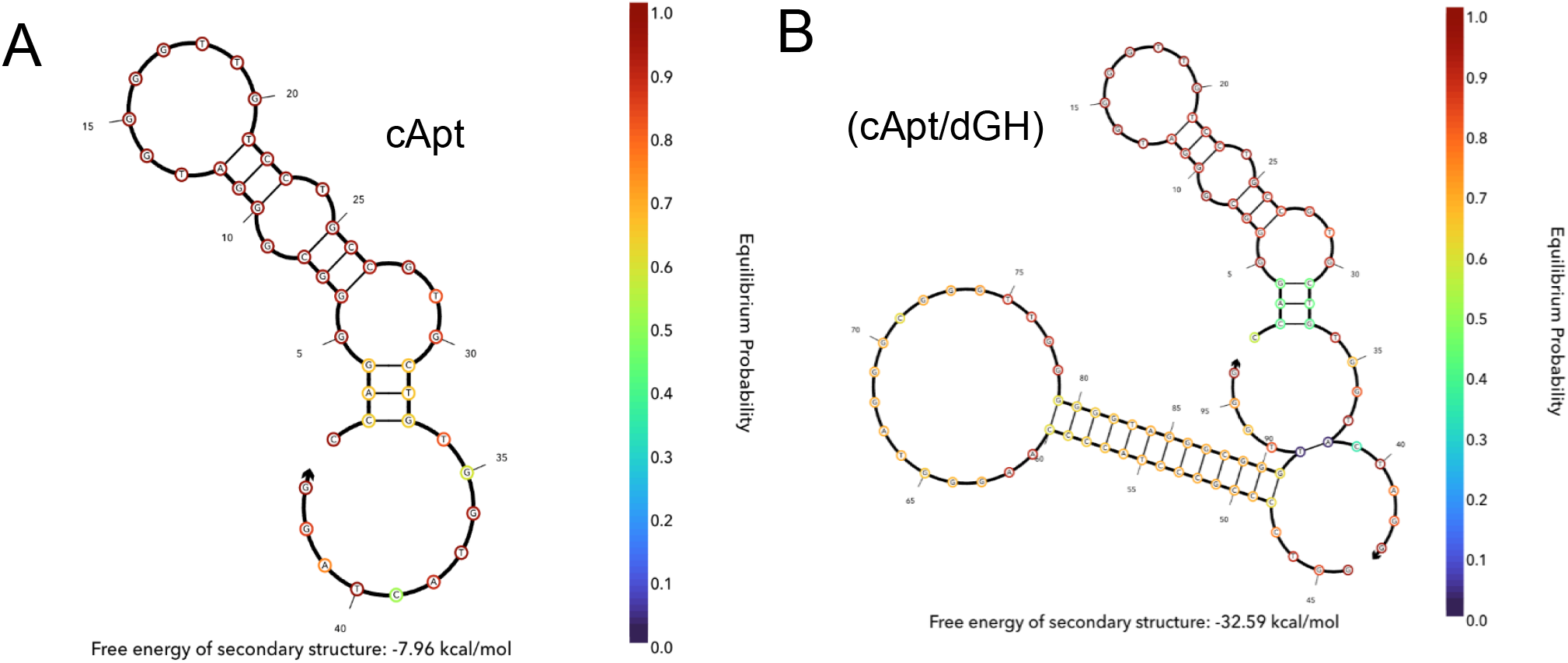
Modelling hybridization of the initiator sequence (cApt) to the g-quadruplex dimer hairpin(dGH) by analyzing Gibbs free energy in 150mM Na+ and 2mM Mg at 21°C using NUPACK software. **A**. Gibbs free energy of cApt alone is -7.96 kcal/mol **B**. Gibb’s free energy of cApt/dGH complex showing a significant drop to -32.59 kcal/mol showing thermodynamic stability confirming high hybridization probability.

Gel electrophoresis was used to verify the affinity of PDGF B Apt to PDGF BB and validate opening of dGH as predicted in ***Figure 3***. To demonstrate the affinity of the PDGF B Apt to PDGF BB protein, an electrophoretic mobility shift assay was performed where the PDGF B aptamer was titrated with four concentrations of protein (1, 3, 5 and 10µM) and run on 12% native PAGE as shown in ***Figure 4***. Results show the appearance of bands with a significantly higher molecular weight compared to PDGF B Apt alone indicating the formation of the PDGF B aptamer/PDGF BB complex. The intensity of the bands decreased as the concentration of the protein decreased as shown in ***Figure 4A***. One well (lane 1 in ***Figure 4A***) was run with just PDGF B Apt as a control and no complex band was observed. These results indicate the strong affinity of the selected aptamer to PDGF BB.

**Figure 4:**
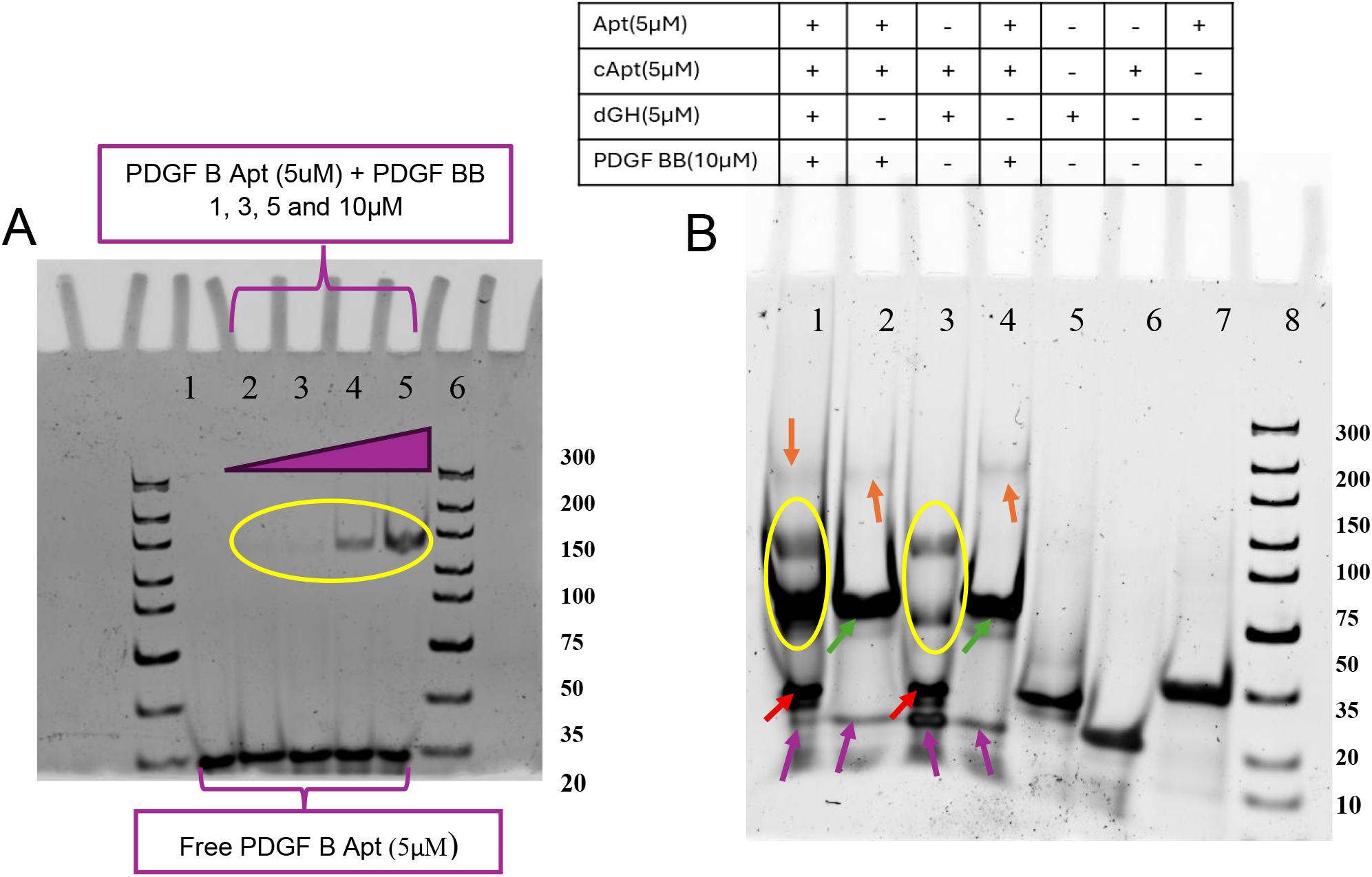
Aptamer affinity and toehold mediated strand displacement assay. **A**. Electrophoretic mobility shift assay where PDGF B aptamer (PDGF B Apt) was incubated with four concentrations of PDGF BB(complexes within yellow circle) lane 1:PDGF B Apt, lane 2: Apt+1 µM PDGF BB, lane 3: Apt+3 µM PDGF BB, lane 4: Apt+5 µM PDGF BB, lane 5: Apt+10 µM PDGF BB, lane 6: Ultra-low range DNA ladder (10-300bp) **B**. Interactions showing lane 1: PDGF B Apt+cApt+PDGF BB+S5, dimeric g-quadruplex resolved as 2 bands within the yellow circle with an overlay of the PDGF B Apt/cApt complex residue, orange arrow shows PDGF B Apt/PDGF BB complex band, purple arrow is cApt residue and red arrow is dGH residue, lane 2: PDGF B Apt +cApt+PDGF BB, green arrow is PDGF B Apt/cApt, orange arrow is PDGF B Apt/PDGF BB complex, purple arrow is displaced cApt, lane 3:cApt+dGH, dGH within the yellow circle, red arrow dGH residue and purple arrow cApt residue lane 4: PDGF B Apt +cApt+PDGF BB, green arrow is PDGF B Apt/cApt, orange arrow is PDGF B Apt/PDGF BB complex, purple arrow is displaced cApt lane 5: dGH, lane 6: cApt, lane 7: PDGF B Apt and lane 8:Ultra-low range DNA ladder (10-300bp).

Binding and hybridization interactions within the TMSD assay were studied with gel electrophoresis. Controls for each sequence were compared with each interaction step within the assay. 12% native PAGE was used to determine the approximate molecular weight of each component, PDGF B Apt, cApt, the initiator sequence with a 4-nucleotide toehold region, and dGH the dimeric g-quadruplex hairpin complementary to cApt all at a concentration of 5µM. Controls for each sequence were run using PAGE. Results from interaction experiment showed that the PDGF B Apt had a slightly higher molecular weight when compared to the dimeric g-quadruplex hairpin (dGH) and initiator sequence (cApt) and (***Figure 4B***). PDGF B Apt/cApt hybridized successfully, with PDGF B Apt/cApt/PDGF BB showing a faint band indicating the binding event between the aptamer and PDGF BB forming a PDGF B Apt/PDGF BB complex. Formation of the complex signaled the successful displacement of cApt from the hybridized PDGF B Apt/cApt complex after protein binding which showed a band on the gel (***Figure 4B***). The displacement of cApt was validated twice in lanes 2 and 4 to demonstrate repeatability of PDGF BB binding to PDGF B Apt. Furthermore, the full assay yields the dimeric g-quadruplex structure shown by the two bands in lane 1 ***Figure 4B***, with the PDGF B aptamer/PDGF BB complex clearly visible.

### Optimization of TMSD assay parameters

To ensure the TMSD assay performs with minimum background signal interference, fluorescence intensity was optimized by studying the effect of DNA concentrations, incubation time and temperature were investigated as shown in ***Figure 5***. Background fluorescence from PDGF B Apt, cApt, and PDGF BB using a concentration of 40nM was compared with the full assay (PDGF B Apt/cApt/PDGF BB/dGH) as shown in ***Figure 5***.

**Figure 5:**
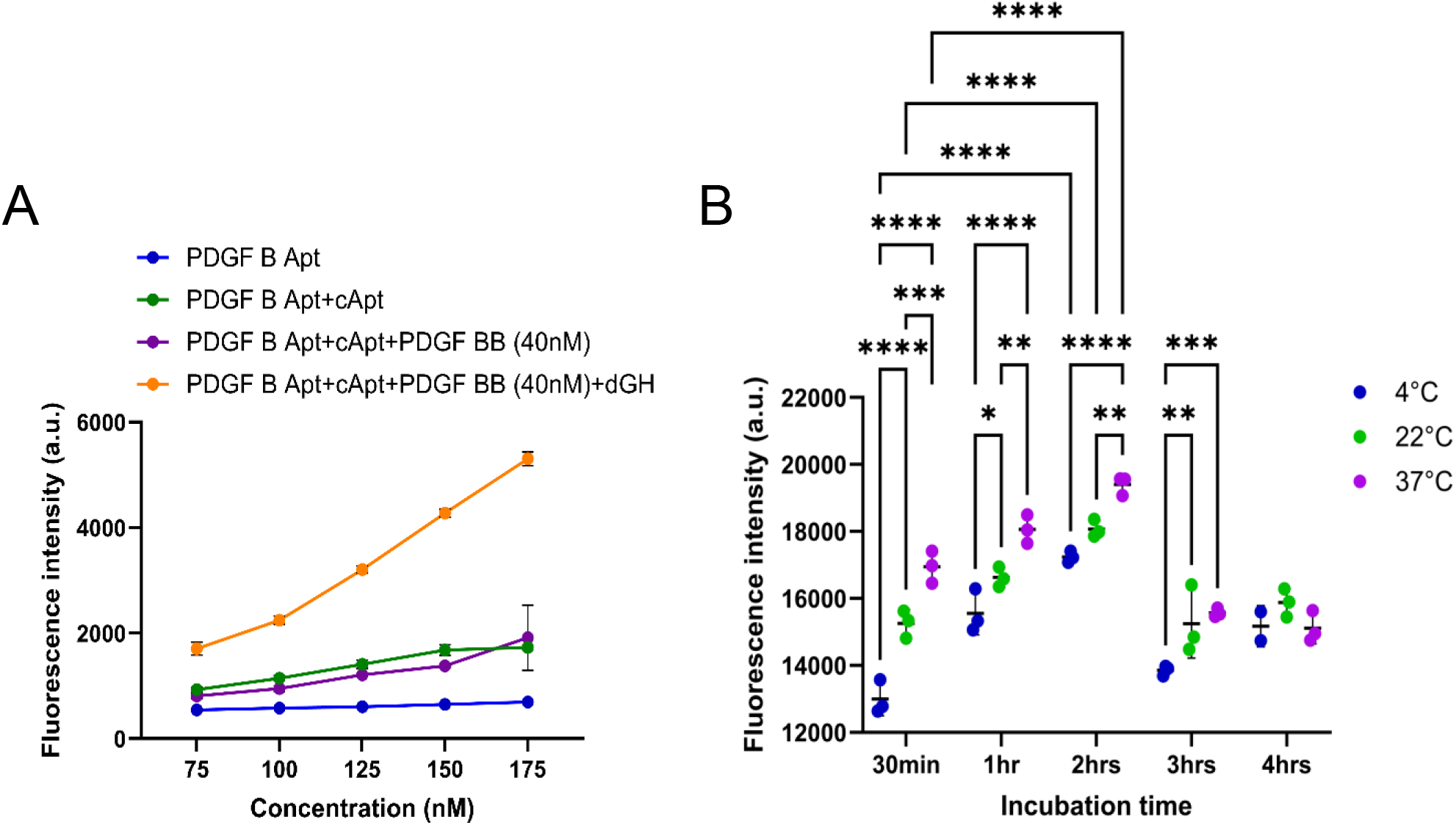
Study on the effect of DNA concentrations, incubation temperature and time on fluorescence intensity. **A**. Fluorescence intensity vs DNA concentration for each step of the assay PDGF B Apt, PDGF B Ap t+ cApt, PDGF B Apt+cApt+PDGF BB and PDGF B Apt+cApt+PDGF BB+dGH **B**. Effect of temperature at 3 levels (4, 22 and 37°C), incubation time at 5 levels (30 minutes, 1, 2, 3 and 4 hours) on fluorescence intensity. *p<=0.05, **p<=0.01, ***p<=0.001, ****p<=0.0001.

Concentrations ranging from 75 to 175nM were prepared with n=3 discrete samples for PDGF B Apt alone, PDGF Apt/cApt, PDGF B Apt/cApt/PDGF BB and PDGF B Apt/cApt/PDGF BB/dGH. Results showed negligible change with the fluorescence intensity with PDGF B Apt alone. With PDGF Apt/cApt and PDGF B Apt/cApt/PDGF BB, there was a slight increase in fluorescence intensity as the concentration increased but tapered off around 150nM for PDGF Apt/cApt with PDGF B Apt/cApt/PDGF BB showing a slight increase. The full assay was observed to steadily increase linearly with an increase in concentration. It is tempting to select the highest concentration for the assay components, however care must be taken with TMSD assays as increasing strand concentrations can skew actual kinetics by limiting rate behavior and obscuring dependence on toehold length and sequence design^27^. Furthermore, high oligonucleotide concentrations increase off-target interactions, and unintended strand displacement, conversely low DNA concentrations slow kinetics, suffer from poor signal to noise^28^. With this background 150nM was chosen as the concentration for the TMSD oligonucleotides since it represented the inflection point where the fluorescence signal strength to linearity changed for background fluorescence with a satisfactory signal to noise ratio for the fluorescence intensity response (***Figure 5A***). A 3x5 factorial design experiment was conducted to understand the relationship between temperature and incubation time on fluorescence intensity. The temperature ranges were selected such that the maximum temperature used would not cause denaturation of PDGF BB protein. Several human and mesophilic proteins start to denature around 40°C - 60°C and are substantially unfolded by 70°C to 95°C^29^. Based on this background the maximum temperature used in the experiment was 37°C. Results show that for each time point there are statistically significant differences with fluorescence intensity for all three temperatures under investigation (***Figure 5B)***. The interaction effect between incubation time and temperature was very significant (p < 0.0001) meaning the effect of temperature levels on the fluorescence intensity, was heavily dependent on the incubation time. Fluorescence intensity reached a threshold at about 2 hours and then dropped at 3 hours (***Figure 5B)***. At 4 hours the fluorescence intensity at each temperature level was not significantly different from 3 hours, and there was no difference between the fluorescence intensity at 4°C, 22°C or 37°C at 4 hours. ThT fluorescence is known to enhance upon binding to amyloid fibrils due to restricted rotation, however, for longer incubation times, higher binding densities occur which causes ThT molecules to be closely packed resulting in self-quenching and reduced emission intensity nonlinearly^30^. With limited information in the literature on the effect of incubation time on fluorescence intensity of ThT for dimeric g-quadruplex scaffolds, this phenomenon could be used to explain why a decrease in fluorescence intensity was observed^31^. In our case, binding of ThT to the dimeric g-quadruplex scaffold via end stacking at the interfaces between G-tetrads and groove binding restricts rotor motion of ThT leading to enhanced fluorescence intensity. However, with prolonged incubation, ThT may saturate the high affinity dimeric g-quadruplex sites and redistribute into weaker external/stacking sites, aggregates on DNA or within the bulk causing a drop in fluorescence intensity. For the purposes of biosensing, short incubation times translate into quicker results read out. Based on the data, incubation at room temperature (22°C) for 30 minutes is ideal and yields sufficient fluorescence intensity for the purposes of the application.

### 3D printed hybrid paper-polymer vertical flow device

3D printing using polypropylene created robust hydrophobic barriers on chromatography paper enabling efficient containment of fluid within the reaction pads (**Figure 6*A&C***). Each reaction pad was designed to have a diameter of 4mm (**Figure 6*A***). Three layers of hybrid paper-polymer devices were integrated within a 3D printed casing with pressure contraptions that clamped the paper-polymer devices in place, aligning reaction pads on each layer (**Figure 6*B&D***), to allow seamless wicking from one layer to another, addressing a common challenge with poor adhesion and inefficient wicking between layers of vertical flow devices. The layers were divided into the sensing layer (2), reagent layer (3) and the waste layer (4) (***Figure 7***). The waste layer was responsible for absorbing excess sample volume, acting like a ‘capillary sink’ by preventing back flow and sustaining unidirectional vertical flow.

**Figure 6:**
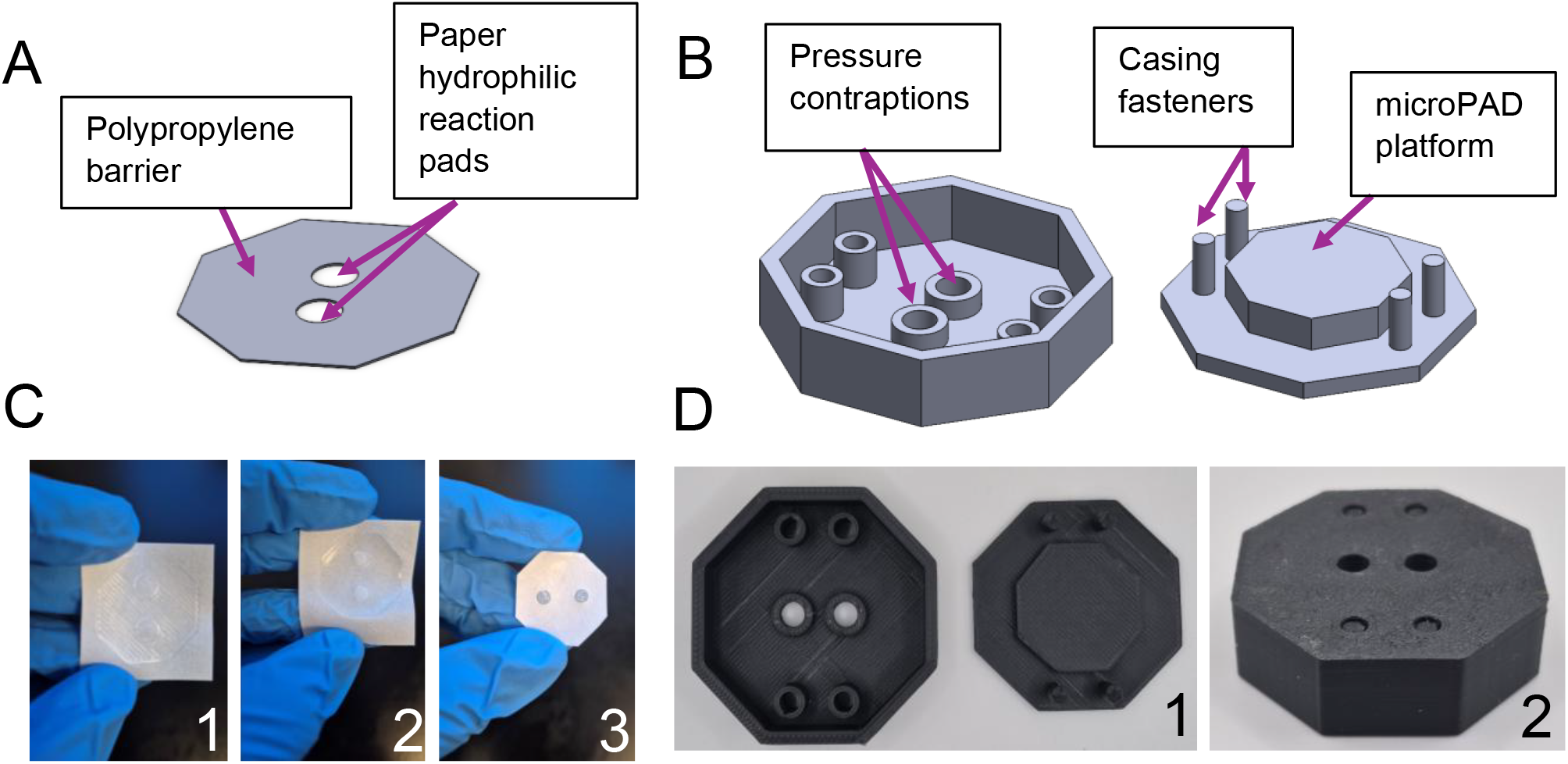
3D printed hybrid paper-polymer vertical flow device design and fabrication. **A**. hybrid paper-polymer microPAD design with hydrophilic paper reaction pads, **B**. device casing showing pressure contraptions to allow seamless wicking between layers **C**. 1: As fabricated 3D printed polypropylene on paper 2: device after thermal curing 3: device with solution deposited, **D**. 1: 3D printed device casing and 2: Fully assembled device.

**Figure 7:**
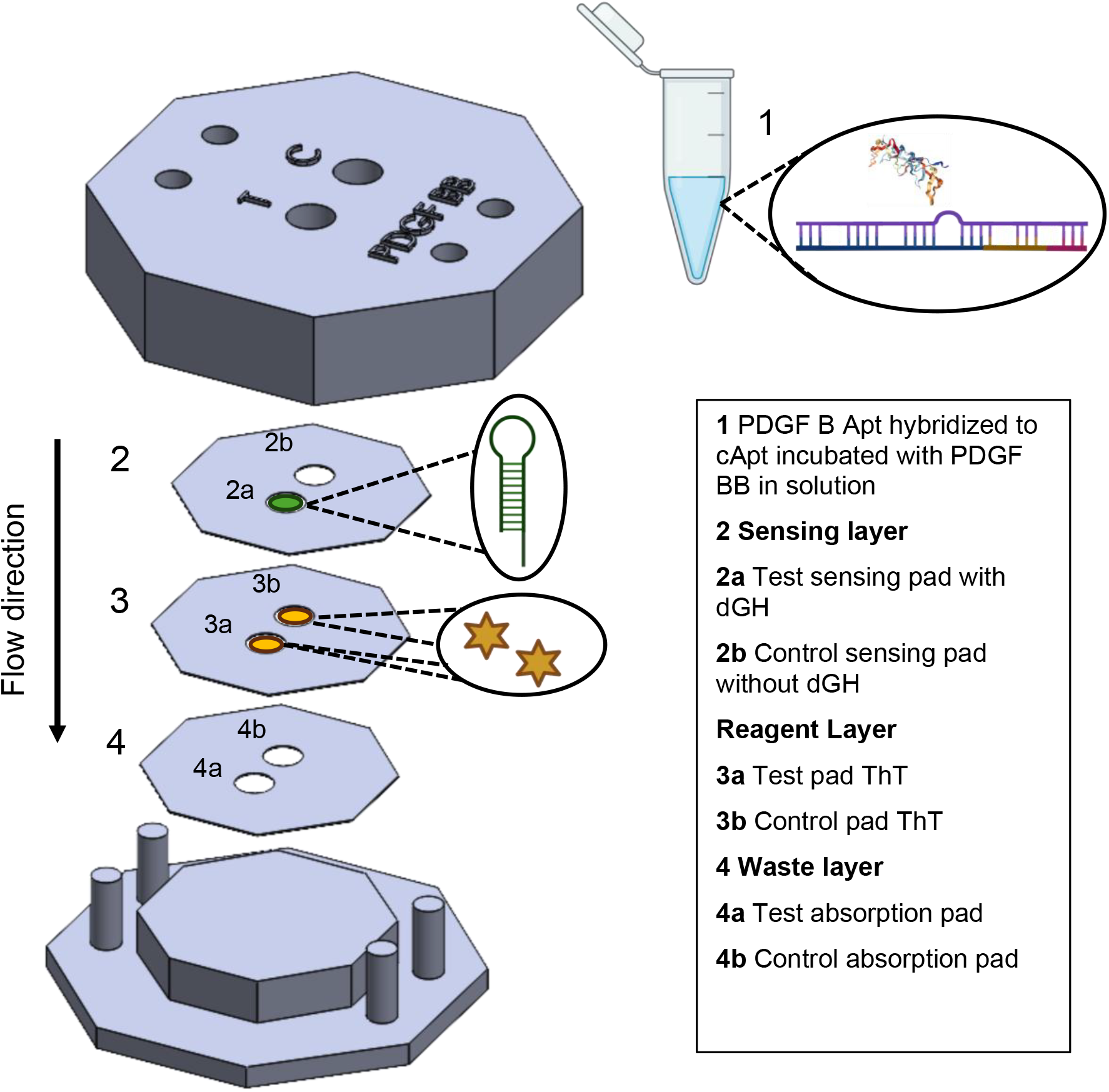
TMSD assay validated on 3D printed Hybrid paper-polymer vertical flow device showing implementation partly in solution. (PDGF B aptamer/cApt incubated with PDGF BB for 30 minutes then deposited on test and control reaction pad enabling wicking from 2 to 4 and subsequently reading the result on layer 2 based on the interaction between layers 2 and 3.

The TMSD assay was first validated in solution and then on the 3D HPVF device (***Figure S1***). From the results in solution, there was a clear distinction between background fluorescence and the formation of the dimeric g-quadruplex structure, indicating the presence of PDGF BB (***Figure S1***). Validation of the TMSD assay on the 3D HPVF device was implemented with first the displacement reaction in solution, then subsequently the hybridization reaction between cApt and dGH on paper (***Figure 7***). The device was designed such that the control reaction pad did not have dGH (2b), therefore the control fluorescence comprised of the background signal (PDGF B Apt/cApt/PDGF BB) without dGH. The PDGF B Apt/cApt was hybridized to form a duplex prior to incubation with PDGF BB protein for 30 minutes at room temperature (22°C). During incubation, the binding event between the PDGF B Apt and PDGF BB triggered the liberation of cApt. A small volume of 10µl of the solution was pipetted onto each sensing pad (test and control) and the solution was allowed to wick through to the reagent layer where on the control pad (3b) cApt and residues from the PDGF B Apt/cApt duplex binds to ThT increasing fluorescence intensity by intercalating between base pairs of the duplex and external binding to phosphate groups (***Figure 7***) . On the test sensing pad, cApt initiates hybridization with dGH, opening the hairpin, fully exposing the nucleotides responsible for forming the dimeric g-quadruplex scaffold. Incubation is at room temperature (22

°C) and the time between the deposition of the sample onto the testing layer to result readout is 30 minutes for a total of 1 hour.

For concentrations of PDGF BB ranging from 10 to 100pM, statistical significance between the fluorescence intensity of the test and control sensing pads was clearly observed (***Figure S2***). In comparison to the results obtained in solution (***Figure S1***), variations with the statistical significance between the test and control regions were observed with 20pM (p < 0.05), 10, 40 and 60pM (p<0.001) and 100pM (p< 0.001) which can be attributed to the complexity of the porous structure (***Figure 8A***). Opening dGH on paper underwent reduced degrees of freedom during hybridization with cApt and during the formation of the dimeric g-quadruplex scaffold, which can be an important factor causing non-linear fluorescence intensity. The approach and goal of this study is indirect (on/off) biosensing of PDGF BB using presence of the dimeric g-quadruplex as an indicator of protein within the sample, therefore distinguishing between background noise and the main signal is key, which was achieved.

**Figure 8:**
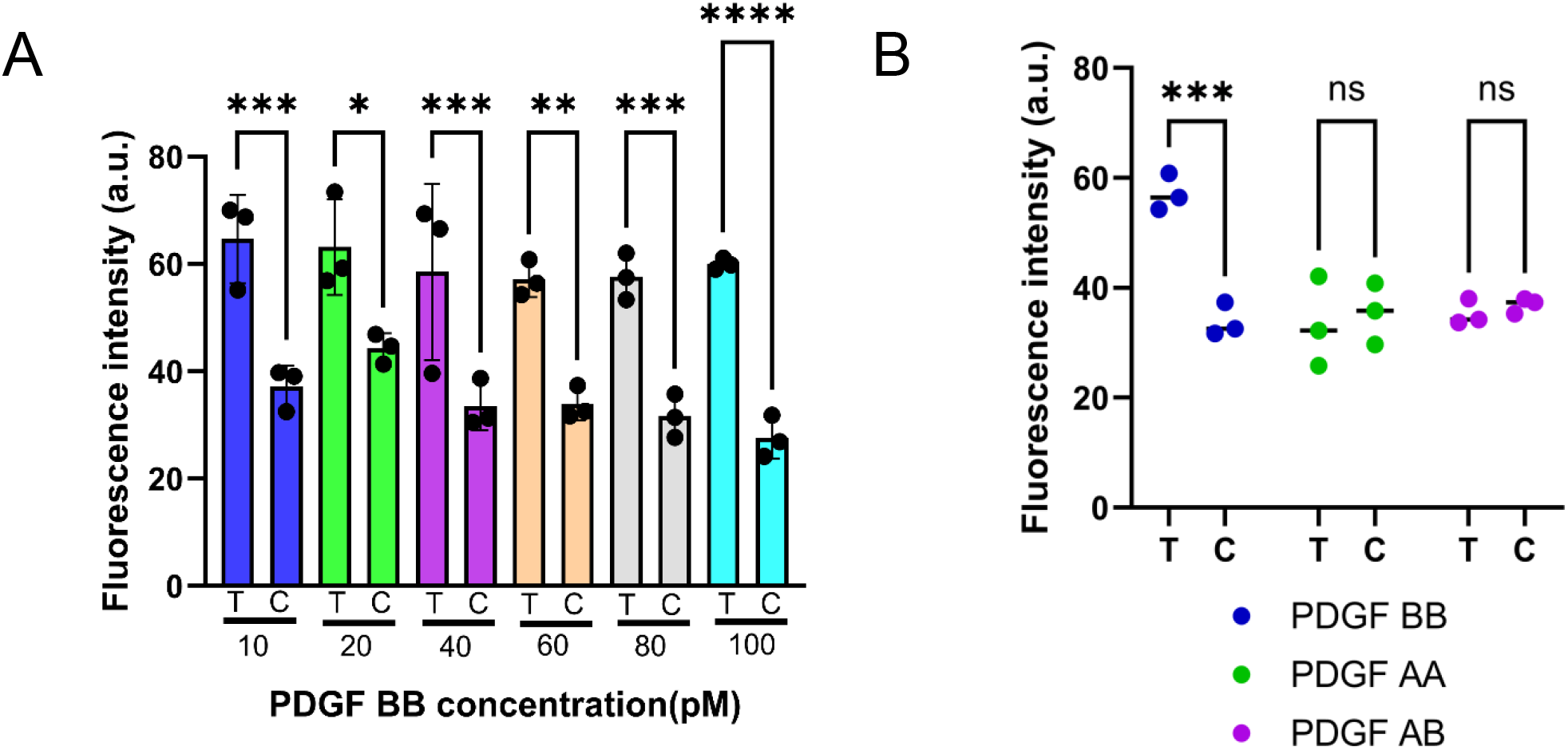
Sensitivity and specificity for 3D HPVF device. **A**. Fluorescence intensity comparisons between the test and control reaction pads of the 3D HPVF device for PDGF BB concentrations ranging from 10pM to 100pM. **B**. Specificity data comparing PDGF BB (60pM) with PDGF AB (1nM) and PDGF AA (1nM). *p<=0.05, ***p<=0.001, ****p<=0.0001.

The 3D HPVF exhibited selectivity for PDGF BB when compared with closely similar proteins PDGF AA and PDGF AB (***Figure S3***), where no significant difference was observed between the test and control of both competing analytes as shown in ***Figure 8B***.

## Conclusion

This work demonstrates the implementation of TMSD on a physical substrate using a 3D HPVF device for qualitative on/off biosensing of PDGF BB. Comprehensive experiments on gel and in solution validated the assay design preparation for implementation on paper. Thorough investigation on the effects of concentration enabled enhancement of signal to noise ratio for the assay. Studies on the effects of temperature and incubation time showed statistical significance between the two parameters and a deeper understanding of the behaviour of fluorescence intensity for dimeric g-quadruplex scaffolds/ThT complexes at different temperature levels over time. 3D printed PP barriers were pivotal in ensuring containment of fluids during the assay run, creating an optimal environment for hybridization and secondary structure formation. The 3D HPVF device achieved an LoD of 10pM showing its potential for sensitive and specific screening of PDGF BB for early onset preeclampsia. The modularity of the design using DNA logic gates enables this platform to be further expanded for multiplexed detection of analytes. Future work will investigate further miniaturization of reaction pads to reduce on variations in fluorescence intensity distribution on paper.

## Acknowledgements

This work was supported by a Natural Sciences and Engineering Research Council of Canada (NSERC) Vanier Canada Graduate Scholarship awarded to P.N., NSERC Discovery Grant (RGPIN-2020-04559) of K.K., Canada Foundation for Innovation John R. Evans Leaders Opportunity Fund of K.K. and R.P., University of Calgary Research Excellence Chair Award and NSERC-AI Advance grant and A-MEDICO grant of R.P.

## Data Availability

The data that support the findings of this study have been included in supplementary information and are available upon reasonable request.

## Conflict of interest

The authors declare no conflict of interest.

## Notes

### Competing Interest Statement

The authors have declared no competing interest.

