## Supplementary Information for "Label-free toehold mediated strand displacement on 3D printed hybrid paper-polymer platform for protein sensing"

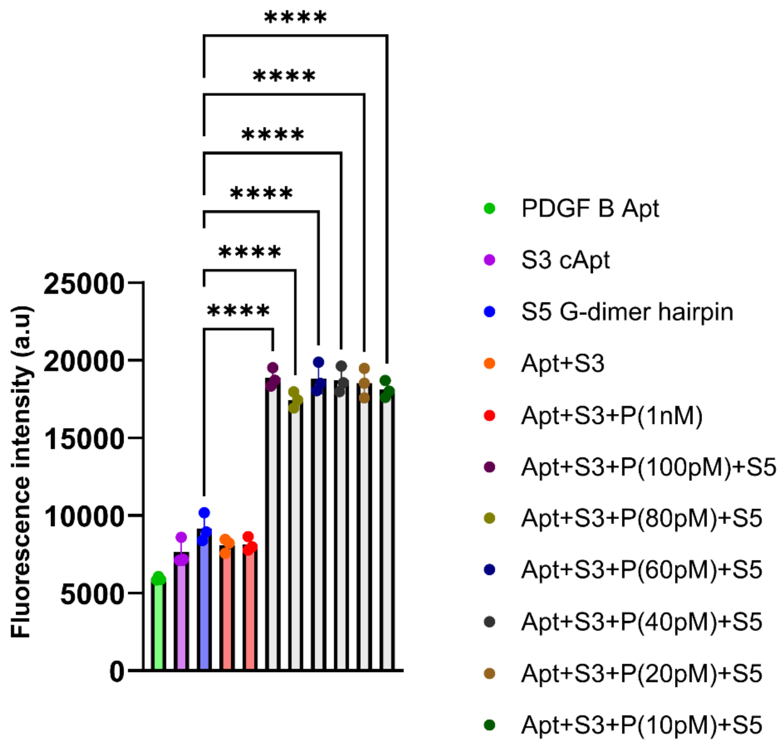


***Figure S1: Validation of the toehold strand displacement assay in solution.*** *Controls are PDGF B aptamer, CAPT complementary to PDGF B aptamer, S5 G-dimer hairpin, Apt+CAPT+1nM to determine background fluorescence with a concentration slightly higher than 100pM and the full assay incubated with concentrations of PDGF BB from 100pM to 10pM. *p<=0.05, **p<=0.01, ***p<=0.001, ****p<=0.0001.*

***Table S1****: Sequences used in the toehold mediated strand displacement assay for PDGF BB detection*

| | |
|---|---|
| PDGF B Apt | 5' GTC CGC TA AA CAG GCT ACG GCA CGA CGT AGA GCA TCA CCA TGA TCC TG 3' |
| cApt | **3' CCAG** GGG CGGG AT GGG GTT GTC CTG CCG TGC TGT GGT ACT AGG 5' |
| dGH | 5' **GGTC** CCC GCCC TA CCC CAA GGG TAG GGC GGG *TTG GGG* GG*TAGG GCG GG*T TGG G 3' |

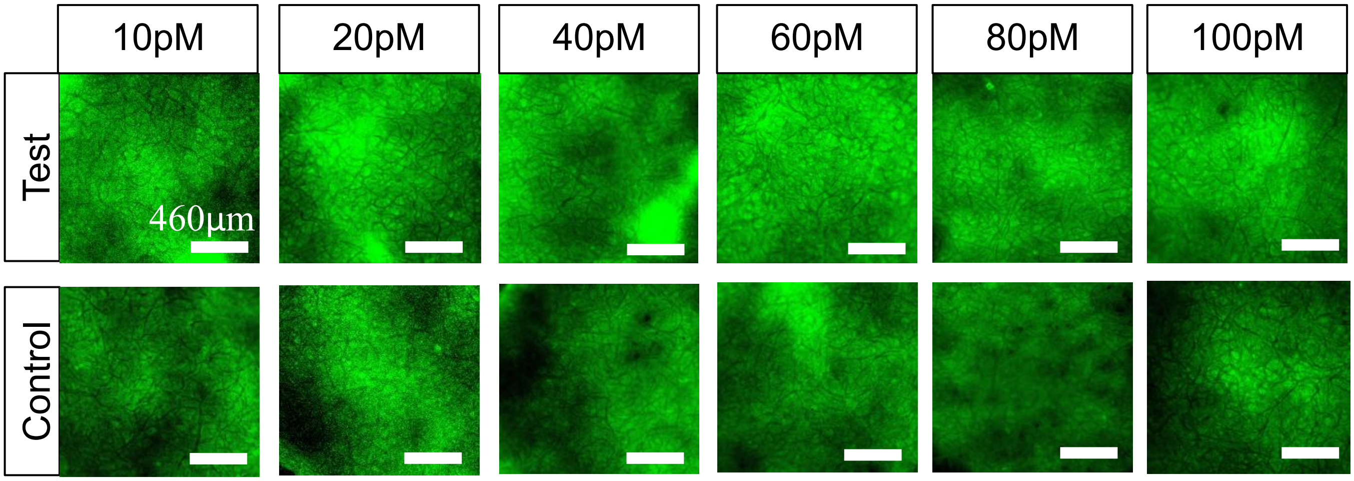


***Figure S2: Fluorescence intensity images for sensitivity of toehold strand displacement assay on 3D hybrid paper-polymer device.*** *10pM to 100pM of PDGF BB was run on the device for 30 minutes incubation time, the control was PDGF Apt/cApt/ PDGF BB ( 10 to 100pM).*

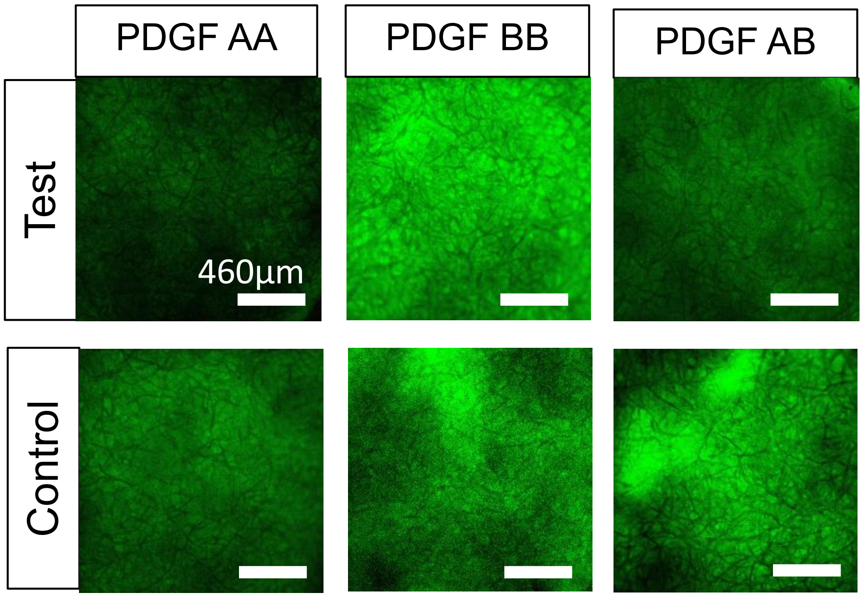


***Figure S3: Fluorescence intensity images for specificity of toehold strand displacement assay on 3D hybrid paper-polymer device.*** *60pM of PDGF BB was compared with 1nM of PDGF AB and PDGF AA. The controls were PDGF B Apt/cApt/PDGF AA, PDGF B Apt/cApt/PDGF BB and PDGF/cApt/PDGF AB.*
